# Oral immunization of goats and foxes with a recombinant NDV vectored rabies vaccine

**DOI:** 10.1101/2023.09.06.556473

**Authors:** Magdalena Murr, Conrad Freuling, David Pérez-Bravo, Christian Grund, Thomas C. Mettenleiter, Angela Römer-Oberdörfer, Thomas Müller, Stefan Finke

## Abstract

Vaccination of the reservoir species is a key component in the global fight against rabies. For wildlife reservoir species and hard to reach spillover species (e. g. ruminant farm animals), oral vaccination is the only solution. In search for a novel potent and safe oral rabies vaccine, we generated a recombinant vector virus based on lentogenic Newcastle disease virus (NDV) strain Clone 30 that expresses the glycoprotein G of rabies virus (RABV) vaccine strain SAD L16 (rNDV_G_RABV_). Transgene expression and virus replication was verified in avian and mammalian cells.

To test immunogenicity and viral shedding, in a proof-of-concept study six goats and foxes, representing herbivore and carnivore species susceptible to rabies, each received a single dose of rNDV_G_RABV_ (10^8.5^ TCID_50_/animal) by direct oral application. For comparison, three animals received the similar dose of the empty viral vector (rNDV). All animals remained clinically inconspicuous during the trial. Viral RNA could be isolated from oral and nasal swabs until four (goats) or seven days (foxes) post vaccination, while infectious NDV could not be re-isolated. After four weeks, three out of six rNDV_G_RABV_ vaccinated foxes developed RABV binding and virus neutralizing antibodies. Five out of six rNDV_G_RABV_ vaccinated goats displayed RABV G specific antibodies either detected by ELISA or RFFIT. Additionally, NDV and RABV specific T cell activity was demonstrated in some of the vaccinated animals by detecting antigen specific interferon γ secretion in lymphocytes isolated from pharyngeal lymph nodes. In conclusion, the NDV vectored rabies vaccine rNDV_G_RABV_ was safe and immunogenic after a single oral application in goats and foxes, and highlight the potential of NDV as vector for oral vaccines in mammals.

**Author Summary:** Oral vaccination of rabies reservoir and spill-over species is the key to control the disease and prevent human rabies. In the past, baits containing live-attenuated rabies vaccines decreased significantly carnivore-mediated rabies in Central and Western Europe as well as North America. However, certain susceptible species are refractory to the oral immunization using so far licensed vaccines. Our vector vaccine based on avian Newcastle disease virus (NDV) has the potential to contribute to the targeted rabies eradication as it was safe and immunogenic after oral administration in goats and foxes. A single vaccine application elicited a rabies virus (RABV) specific systemic humoral immune response in the majority of the vaccinated animals as well as RABV specific T cells in some of the vaccinated animals. NDV can be manufactured at low-cost using already existing infrastructure of influenza vaccines, opening new possibilities especially for middle- and low-income countries that suffer under the economically burden of rabies.

## Introduction

Rabies is a viral zoonotic infectious disease of the central nervous system with fatality rates of almost 100 % once symptomatic and is caused by rabies virus (RABV, species: *rabies lyssavirus*), a neurotropic virus, belonging to the genus lyssavirus within the family *Rhabdoviridae.* Combating the disease in wild life reservoir species can only be facilitated by oral vaccination using live, replication-competent viruses. From classical attenuated to biotechnological approaches, various oral rabies virus constructs have been developed for use in wildlife and dogs. Attenuated RABV vaccines include 1^st^ to 3^rd^ generation vaccine virus constructs, which almost all descend from the same progenitor virus, designated as “Street Alabama Dufferin” (SAD), isolated from a rabid dog in Alabama, USA, in 1935 (1). While 1^st^ generation attenuated rabies vaccines have been obtained by serial passaging, plaque purification or clonal selection, 2^nd^ and 3^rd^ generation were generated by anti-glycoprotein (G) monoclonal antibody driven selection and targeted site-directed mutagenesis at important antigen sites of the former, respectively (2). The different development stages thereby reflect improvements in residual pathogenicity, with the 3^rd^ generation vaccine virus constructs showing the highest safety profile (3). Although many of the constructs developed are based on proof-of-concept studies and do not make it to market, to date, attenuated rabies vaccines are still the most commonly used vaccines in oral rabies vaccination (ORV) programs worldwide and have been instrumental in the elimination of rabies in foxes in Western Europe and North America.

Biotechnology-derived or genetically engineered oral rabies vaccines include both 3^rd^ generation attenuated RABV and vector virus-based vaccines. As for the latter, various vector viruses have been constructed for the expression of RABV glycoprotein (G) encompassing several virus genera and families, among them vaccinia virus (VACV), human adenovirus 5 (hAdV) and canine adenovirus 2 (cAdV), pseudorabies virus (PrV), parainfluenza virus 5 (PIV5), and Newcastle disease virus (NDV) of which only recombinant hAdV (ONRAB) and VACV (Raboral V-RG) are licensed in certain countries for oral rabies vaccination of wildlife so far (4–8).

Oral rabies vaccines elicit an immune response via the oral cavity. Recent studies indicated that the palatine tonsils of meso-carnivores are a major site of vaccine virus uptake. However, different species depicted a divergent responsiveness after oral immunization, ranging from very sensitive to rather refractory, as shown by the titer required to trigger an immune response. While responsive species including foxes (*Vulpes Vulpes*), raccoon dogs (*Nyctereutes procyonoides*) and mongooses (*Herpestes auropuncatus*) can already be vaccinated with relatively low minimum effective vaccine virus titers, skunks and raccoons seem to be rather refractory to oral rabies vaccination, even when high virus titers were administered regardless whether attenuated or biotechnology derived vaccines were used (9–14). The same is probably true for the Greater Kudu (*Tragelaphus strepsiceros*), a species important for wildlife farming in Namibia for which oral immunization is the only viable preventive measure to avoid substantial numbers of death due to rabies (15).

Therefore, the development of novel, highly safe, environmentally robust and cost-effective vaccines that are immunogenic in various animal species by the oral route would be desirable.

We here evaluated Newcastle disease virus (NDV), a single stranded negative sense RNA virus (species: *Avian orthoavulavirus 1*) belonging to the genus *Orthoavulavirus* of the family *Paramyxoviridae* (16, 17) as a potential oral vaccine candidate. Virulent (velogenic and mesogenic) NDV strains cause Newcastle disease (ND) in avian species, a notifiable epizootic with high mortality rates in naïve populations which is endemic in various countries in Africa, Asia, Central and South America (18, 19). Lentogenic (low-virulent) NDV strains are naturally attenuated in birds and are used as live-attenuated vaccines to control and prevent ND in poultry, but also as backbone for recombinant vector vaccines in poultry and mammals (20).

Replication-competent lentogenic NDV vectored rabies vaccines were shown to be safe and highly immunogenic by inducing a strong long-lasting humoral and cell mediated immune response in mice, dogs and cats after repeated intramuscular (i. m.) inoculation (8, 21, 22). However, their potential in eliciting an immune response after oral application has never been explored in detail. Therefore, we set out to generate an NDV based recombinant vector virus expressing the RABV G of vaccine strain SAD L16 and test its immunogenicity after a single oral application. In this proof-of-concept study we used goats and foxes as model species for herbivores (i.e. Kudu) and carnivore rabies reservoir species.

## Materials and methods

### Cells and viruses

BSR-T7 cells (baby hamster kidney cells, BHK 21, clone BSR-T7/5; CCLV-RIE 582) (23), which stably express phage T7 polymerase, were used for recovery of recombinant NDV, and they were maintained and grown in Glasgow minimal essential medium, supplemented with NaHCO_3_, casein peptone, meat peptone, yeast extract, essential amino acids, and 10 % fetal calf serum (FCS). Geneticin (G418 sulfate; 1mg/mL) was added weekly to the culture to select T7 polymerase positive cells. Primary chicken embryo fibroblasts (CEF) were prepared from 10-day-old specific pathogen free (SPF) embryonated chicken eggs (ECE), purchased from Valo, BioMedia (Osterholz-Scharmbeck, Germany) and incubated at 37 °C with 55 % humidity. CEF, DF-1 cells (permanent chicken embryo fibroblasts, CRL-12203), QM9 cells (quail muscle cells, clone 9, CCLV-RIE 466), BHK-21 (baby hamster kidney cells, CCL-10), and MDBK (Madin Darby bovine kidney cells, CCLV-1193) were used for virus characterization, maintained and grown in minimal essential medium, supplemented with NaHCO_3_, Na-Pyruvate, non-essential amino acids, and 10 % FCS. All cells were incubated at 37 °C with 3 % - 5 % CO_2_.

Recombinant NDV (rNDVGu) based on lentogenic NDV Clone 30 (Genbank Acc. No. Y18898) has been described before and is further on referred to as rNDV (24). RABV strain SAD L16 (25) is a recombinant clone of RABV SAD B19 (Genbank Acc. No. EU877069) and was used for in *in vitro* lymphoycyte re-stimulation. RABV CVS-11 (challenge virus standard-11, ATCC VR 959) (26) was used for virus neutralization assay.

### Construction and generation of recombinant NDV expressing RABV glycoprotein (G)

The open reading frame (orf) encoding RABV SAD L16 G was amplified from pCMV-SADL16 (27) using the Expand High Fidelity^PLUS^ PCR System (Roche Applied Science, Mannheim, Germany) with specific primers PRVGncrNDF (5’-<u>CTA CCG CTT CAC CGA CAA CAG TCC TCA ACC</u> ATG GTT CCT CAG GCT CTC CTG TTT GTA CC-3’) and PRVGncrNDR (5’-<u>CCA ACT CCT TAA GTA TAA TTG ACT CAA</u> TTA CAG TCT GGT CTC ACC CCC ACT CTT GTG-3’). The primers contain parts of the NDV 3’ and 5’ non-coding regions (ncr), derived from the NDV hemagglutinin-neuraminidase (HN) gene (underlined primer sequence parts). The 1.6 kb PCR product was gel-purified using the QIAquick® Gel Extraction Kit (Qiagen, Hilden, Germany) and was subsequently inserted into a cloning vector pUCNDVH5 (28) by Phusion polymerase chain reaction (Finnzymes Phusion®, New England Biolabs®) (29), thereby replacing the H5 insert from pUCNDVH5 with the G orf, resulting in pUCNDV_G_RABV_, in which the G orf is inserted between NDV F and HN genes, flanked by NDV HN ncr. Cloning vector pUCNDV_G_RABV_ and cDNA full-length plasmid pNDVGu were both cleaved with *Not*I and *BsiW*I. The *Not*I-*BsiW*I-fragment of pNDVGu was subsequently replaced with the 5.8 kb gel-purified *Not*I-*BsiW*I-fragment of pUCNDV_G_RABV_, resulting in cDNA full-length plasmid pNDV_G_RABV_.

### Transfection and virus recovery

Infectious recombinant NDV was rescued and propagated as described (30). The full-length plasmid pNDV_G_RABV_ was co-transfected with helper plasmids pCiteNP, pCiteP, and pCiteL into BSR-T7 cells using Lipofectamine®3000 (Invitrogen, Carlsbad, USA) and a DNA to Lipofectamine ratio of 1.0 µg to 1.5 µL, following the manufacturer’s instruction.

### RNA isolation, reverse transcriptase (RT)-PCR, and sequencing

RNA was isolated from allantoic fluid after the first, second and 10^th^ passage in ECE of recombinant NDV_G_RABV_ using the QIAamp® Viral RNA Mini Kit (Qiagen). Genomic regions, encoding the proteolytic cleavage site within the NDV F gene, as well as the region, encoding the inserted RABV G, were transcribed into cDNA and amplified, using the Transcriptor One-Step RT PCR Kit (Roche Applied Science, Mannheim, Germany). Sanger-sequencing (Sequencer 3130 Genetic Analyzer, Applied Biosystems, Foster City, CA, USA) was used to confirm virus identity. Virus stock used for *in vitro*-characterization and *in vivo*-experiments was prepared from allantoic fluid of the second ECE passage.

### Primary antibodies and sera

Mouse monoclonal antibodies (mAb), monospecific rabbit antisera as well as a hyperimmune serum against NDV (HIS α NDV), were used to detect viral proteins by indirect immunofluorescence assay (IFA) or Western blotting (table 1). mAb β-Actin (Sigma-Aldrich, Darmstadt, Germany) was used in Western blot analyses to detect cellular β-actin as loading control.

**Table 1:**
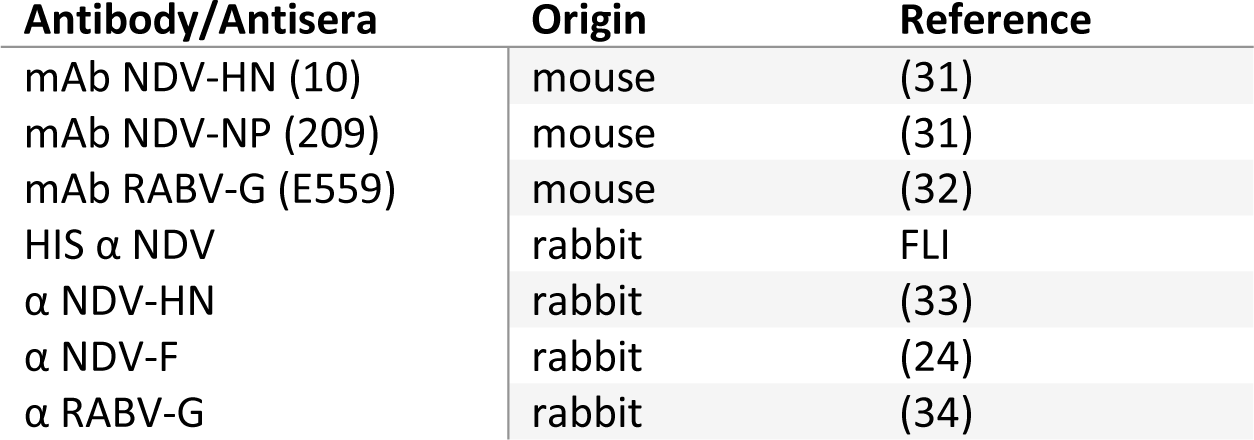
Primary antibodies and sera used for virus *in vitro*-characterization.

### Kinetics of replication

CEF, QM9, MDBK, and BHK-21 cells were infected with rNDV or rNDV_G_RABV_ at a multiplicity of infection (moi) of 0.01. Cell monolayers were washed twice with medium after an adsorption time of 40 min. Cell culture supernatants were harvested 0, 17, 24, 48, and 72 h after infection (p. i.), and examined for the presence of infectious progeny viruses by titration in duplicate on QM9 cells, which were fixed 20 h p. i.. Viral titers (50 % tissue culture infectious dose, TCID_50_) were calculated by IFA using HIS α NDV and Alexa Fluor^TM^ 488 α-rabbit secondary antibody (Invitrogen) in two independent experiments.

### Virus purification

Recombinant virions were concentrated from allantoic fluids by ultracentrifugation on a 60 % sucrose cushion, and purified by ultracentrifugation through a caesium chloride gradient as described before (35).

### Western blot analyses

DF-1, MDBK or BHK-21 cells were infected with rNDV and rNDV_G_RABV_ at an moi of 5, harvested 24 h and 48 h p. i. and lysed in 1x Roti®-Load buffer (Roth, Karlsruhe, Germany). Purified virion solutions were mixed 1:1 with 1x Roti®-Load buffer. All lysates were denatured at 95 °C for 10 min, proteins were separated by sodium dodecyl sulphate polyacrylamide gel electrophoresis (SDS-PAGE), and subsequently transferred to nitrocellulose membranes. Viral proteins were detected by incubation with α RABV-G, α NDV-HN, or α NDV-F. After primary antibody incubation, binding of peroxidase-conjugated species-specific secondary antibodies (Dianova) was detected by chemiluminescence substrate (Thermo Scientific, Rockford, IL, USA) and visualized by ChemiDoc XRS+ (BioRad, Hercules, CA, USA).

### Indirect immunofluorescence assay

QM9 cells were infected with rNDV and rNDV_G_RABV_ at an moi of 0.1, fixed with 4 % paraformaldehyde 24 h p. i., and permeabilized using 0.1% Triton X-100. After blocking of permeabilized and non-permeabilized cells with 5 % bovine serum albumin (BSA) in PBS, they were incubated with α NDV-HN and mAb RABV-G. Binding of primary antibody was visualized using Alexa 488 α-rabbit or 568 α-mouse secondary antibody (Invitrogen). 4’,6-Diamidino-2-phenylindole (DAPI) (Roth, Karlsruhe, Germany) was included in washing steps after binding of secondary antibodies to stain nuclei. Images were taken on a Leica SP5 confocal microscope (Leica Microsystems GmbH, Wetzlar, Germany) with an oil immersion objective (HCX PL APO 63x/1.40–0.60 objective). Sequential z-sections of stained cells were acquired for maximum projection, and images were processed using ImageJ software (36).

### Mean death time (MDT)

Allantoic fluid of rNDV_G_RABV_ was diluted ten-fold serially. Ten 10-day-old SPF ECE each were inoculated with 0.1 mL of each dilution and incubated at 37 °C and 55 % humidity for 168 h. The MDT is defined as the mean time in hours for the minimum lethal dose to kill all embryos. The minimum lethal dose is the highest virus dilution that causes all the embryos inoculated with that dilution to die (18).

## Animal trials

### Animals and housing conditions

The pathotype of rNDV_G_RABV_ was determined in one-day-old SPF-chickens (*Gallus gallus*; mixed sexes), purchased from VALO BioMedia and hatched at the Friedrich-Loeffler-Institute (FLI), Insel Riems. Ten chickens were housed together in a cage and provided with food and water ad libitum. For the oral vaccination studies, a total of nine goats (*Capra aegagrus*, male) and nine red foxes (*Vulpes vulpes*, mixed sexes) were purchased from commercially registered breeders in Germany (Rubkow, Mecklenburg-Western Pomerania) and Poland (Wielichowo, Wielkopolskie), respectively. All animals were clinically healthy upon arrival. Goats were housed together, whereas the foxes were kept in single cages essentially as described before (37). Animals were provided with commercial standard feed once a day, according to individual consumption behaviour and need, except for a weekly fasting day (foxes), and water ad libitum as described before (37).

### Ethical statement

Experimental studies were carried out in biosafety level 2 animal facilities at the FLI, Insel Riems. All animal experiments were approved by the animal welfare committee (Landesamt für Landwirtschaft und Fischerei Mecklenburg-Vorpommern, Thierfelder Straße 18, 18059 Rostock, LALLF MV/TSD/7221.3-1-009/19 and LALLF M-V/TSD/7221.3-1-003/21) and supervised by the commissioner for animal welfare at the FLI, representing the institutional Animal Care and Use Committee (ACUC). The studies were conducted in accordance with national and European regulations, and European guidelines on animal welfare from the Federation of European Laboratory Animal Science Associations (FELASA).

### Intracerebral pathogenicity index (ICPI)

The ICPI was determined in one-day-old SPF chickens, following the standard protocol (38). Briefly, ten one-day-old SPF chickens were inoculated intracerebrally with 100 µL of a 10^-1^ dilution of rNDV_G_RABV_. Mortality and clinical signs were monitored daily for 8 days. Chickens showing severe clinical distress during the experiment were euthanized. Criteria for euthanasia were dyspnea, apathy, somnolence, akinesia, or deficit motor activity, respectively.

### Oral vaccination of goats and foxes

To investigate immunogenicity of rNDV_G_RABV_ *in vivo*, goats and foxes in the test group (n = 6) each were given 3x10^8.5^ TCID_50_/animal of rNDV_G_RABV_ by direct oral application (DOA). The remaining animals were assigned controls (n = 3) receiving 3x10^8^ TCID_50_/animal of parental rNDV by the same route. Clinical signs were monitored daily with a scoring system in the following categories: activity (0–4), posture (0–4), temperature (0–3), fur (0–3), ocular and nasal discharge (0–3), respiration (0–3), feed and water intake (0–3), and defecation (0–2). The criterium for euthanasia was a cumulative score of > 8 within one day. Body temperature of goats and foxes were measured on days of sampling. The body weight of foxes was additionally assessed on the same days.

Blood was taken day 0, 7, 14 and 28 post vaccination (dpv) to monitor humoral immune response. Oral, nasal and rectal swabs were taken 0, 2, 4, 7 and 14 dpv to monitor vaccine virus replication and shedding. For DOA, foxes and sampling foxes were sedated with 0.5 – 1 mL Tiletamin hydrochloride + Zolazepam hydrochloride (Zoletil®, Virbac Arzneimittel GmbH, Bad Oldesloe, Germany) per animal, while goats were restraint using movable cage gates. All animals were euthanized 28 dpv using Zoletil® followed by exsanguination in deep general anesthesia; at necropsy retropharyngeal lymph nodes were taken from every animal for preparation of lymphocytes.

### Virus replication and shedding

Viral RNA from nasal, oral and rectal swabs was isolated automatically (KingFisher 96 Flex, ThermoFisher, Waltham, MA, USA) using the NucleoMag® VET Kit (Machery-Nagel, Düren, Germany). Reverse transcriptase quantitative real-time PCR (RT-qPCR) was used to detect NDV NP specific RNA essentially as described (24). Virus loads determined were expressed as genomic equivalents (GEQ) using calibration curves of defined RNA standards included in each RT-qPCR run. All RT-qPCRs were performed in 12.5 µL volumes using the AgPath-ID RT-PCR Kit (Ambion, Austin, TX, USA) and run on a CFX96 thermocycler machine (Bio-Rad, Feldkirchen, Germany). Cycle threshold (Ct) values of 40 were set as the cut-off, representing a GEQ/mL of 1000.

### Serology

Serum antibody titers against NDV were determined using the hemagglutination inhibition (HI) assay according to the standard protocol, using 4 hemagglutinating units of parental rNDV as antigen (38). HI titers ≥ log_2_ 3 were considered seropositive. NDV neutralizing antibodies were detected in duplicates by virus neutralization assay (VNA) using rNDV as test virus as described (39). The neutralizing antibody titers were defined as the highest dilution which showed virus neutralization in both wells. NDV binding antibodies (ID Screen® Newcastle Disease Competition) were determined using a commercial competitive ELISA (Innovative Diagnostics, Grabels, France). Sera showing an inhibition of 30% or 40% were considered indeterminate or positive, respectively.

RABV neutralizing antibodies in fox and goat sera were measured using a modified fluorescent focus inhibition test (RFFIT) using positive (World Health Organization 2nd International Reference Standard, National Institute of Biological Standards and Controls, Potter’s Bar, UK) and negative controls and RABV strain CVS-11 as test virus as described before (40). Antibody titers ≥ 0.5 IU/mL were considered positive. For the detection of rabies specific binding antibodies, a commercial competitive ELISA (BioPro Rabies ELISA, Prague, Czech Republic) was used. Sera showing an inhibition of 40 % were considered positive (41).

### Isolation of lymphocytes

The pharyngeal lymph nodes of all animals were removed post mortem to isolate lymphocytes. Organs were grinded mechanically using a scissor and subsequently squeezed through a 70 µm cell strainer to separate the cells. Lymphocytes were washed in 1x PBS/EDTA (0.5 %) and resuspended in cell culture medium supplemented with FBS (10%), Penicillin/Streptomycin (100 IU/mL/100 µg/mL; Gibco™) and 2-Mercaptoethanol (50 µM; Gibco™).

### IFN-γ Enzyme-Linked ImmunoSpot (ELISpot) detection assay

NDV and RABV G specific T cell activity was analyzed by detecting antigen specific interferon γ (IFN-γ) production using the canine IFN-γ ELISpotPlus (ALP) Kit for fox lymphocytes and the respective bovine Kit for caprine lymphocytes (Mabtech AB, Nacka Strand, Sweden) following the manufacturer’s instructions. Briefly, lymphocytes isolated from pharyngeal lymph nodes were transferred in duplicates into pre-coated, equilibrated 96-well ELISpot plates (2 × 10^5^ lymphocytes/cavity) and re-stimulated with either β propiolactone inactivated parental rNDV or RABV SAD L16 (5.0 µg/mL). Concanavalin A (3.0 µg/mL; ConA; Sigma-Aldrich-Merck, Darmstadt, Germany) served as positive control antigen and cell culture medium as negative control antigen. Plates were washed after 24 h of stimulation. Secretion of IFN-γ was detected using a biotinylated anti-canine IFN-γ monoclonal antibody, and subsequently streptavidin-ALP and ready-to-use BCIP/NBT-plus substrate. Spots were automatically identified using the vSpot Spectrum ELISpot Reader (AID GmbH, Strassberg, Germany) and counted as Spot Forming Units per one million cells (SFU/10^6^ cells).

### Statistical analysis

Statistically significant differences between viral titers of replication kinetics were analyzed using an unpaired t-test and a significance interval of 95% (α = 0.05).

The Pearson correlation coefficient r was calculated to identify correlation between serological data obtained either with NDV specific assays or RABV specific assay. Significance of correlation was determined applying a significance interval of 95% (α = 0.05). Graphs preparation and statistical analyses were performed using GraphPad Prism Software Version 7.05 (San Diego, CA, USA).

## Results

### Generation of recombinant NDV expressing RABV G

Recombinant NDV_G_RABV_ harbors a transgene encoding the G orf of rabies vaccine strain SAD L16. The foreign gene was inserted into the igr separating F and HN genes of lentogenic NDV strain Clone 30 (rNDV) with appropriate gene start, gene end and non-coding sequences derived from the NDV HN gene (Fig. 1).

**Figure 1.**
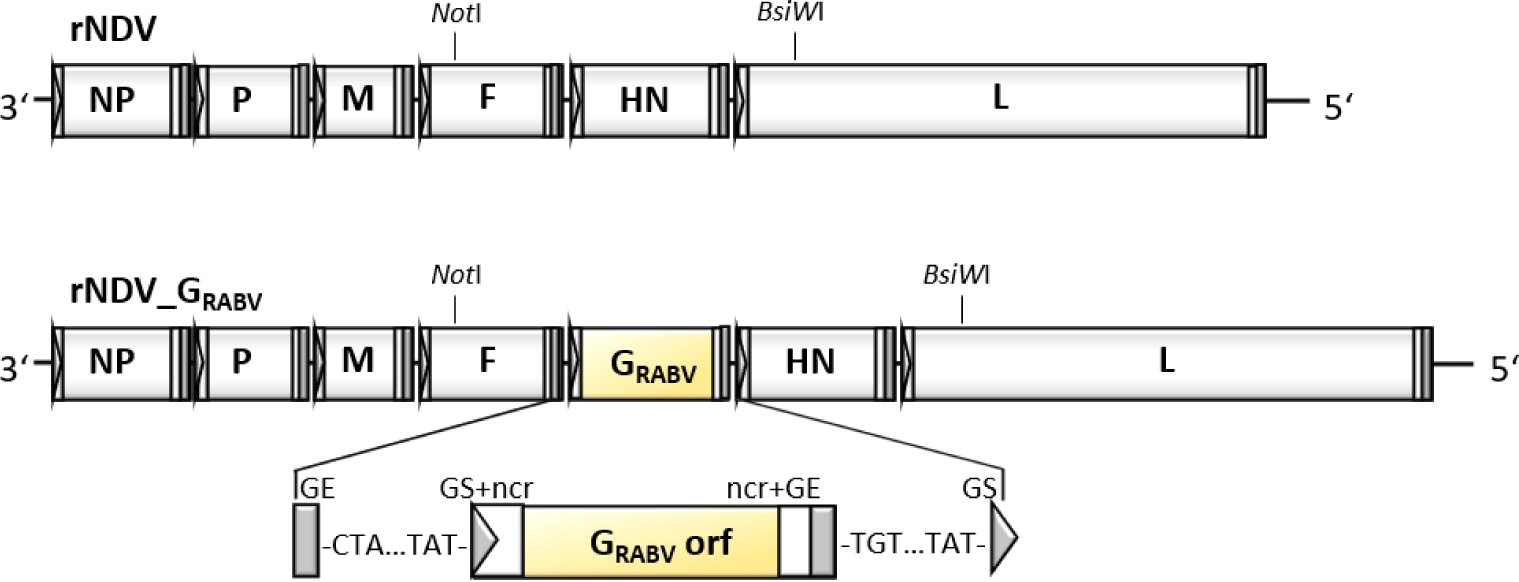
Construction of recombinant Newcastle disease virus (NDV) expressing rabies virus (RABV) glycoprotein (G). The RABV transgene is inserted into the intergenic region (igr) of the NDV fusion protein (F) and the hemagglutinin-neuraminidase protein genes (HN). The open reading frame (orf) is flanked by NDV specific gene start (GS, grey triangle) and gene end sequences (GE, grey rectangle), and by the NDV HN specific non-coding regions (ncr, white rectangles).

Recombinant NDV_G_RABV_ was rescued in BSR T7/5 cells and subsequently propagated in SPF-ECE. After two passages in SPF-ECE, the identity of recombinant NDV_G_RABV_ was confirmed by amplification and sequencing of selected regions of the viral genome. To assess stability of the recombinant virus, NDV_G_RABV_ was passaged ten times in SPF-ECE. RNA of the 10th egg passage was isolated. The genomic regions encoding the proteolytic cleavage site of NDV F, as well as RABV G were sequenced. No alteration was observed in any of the analyzed sequences. These results suggest stability of the RABV G insert over ten SPF-ECE passages.

### Recombinant NDV_G_RABV_ replicates in embryonated chicken eggs and in cell lines originated from different species

Replication efficacy of rNDV_G_RABV_ was investigated in ECE which is the gold standard for NDV propagation and vaccine generation as well as in chosen avian and mammalian cell lines representing NDV and RABV host species. rNDV_G_RABV_ replicated to high titers in ECE which were significantly higher 72 h p. i. compared to parental rNDV (Fig. 2A). Viral titers obtained in cell culture were lower as in ECE. Both viruses reached comparable titers in all investigated cell lines, whereby titers in BHK-21 and MDBK cells were about 1.5-2 log lower 72 h p. i. than in CEF (Fig. 2B-D). Replication in both mammalian cell lines was independent of the supplementation of exogenous trypsin, which is required for lentogenic NDV F cleavage and subsequent production of infectious viral progeny particles, suggesting that the kidney cells produce this type of proteases. In contrast, CEF culture contained trypsin as it is part of the cell preparation. No replication was observed in QM9 cells which lack of adequate activating enzymes (data not shown).

**Figure 2.**
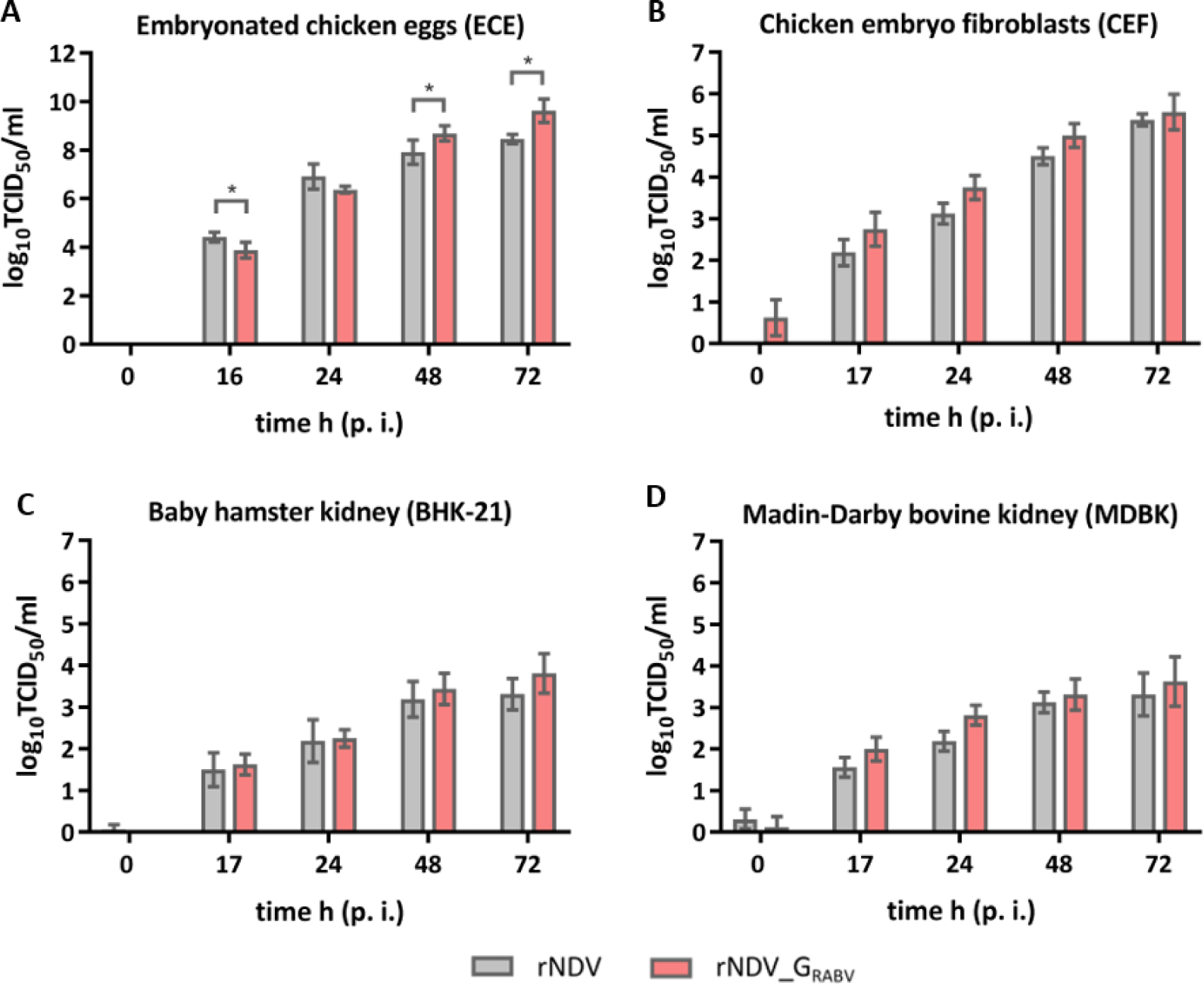
*In vivo-*replication in embryonated chicken eggs and *in vitro* replication in cell lines originated from different species. Embryonated chicken eggs (ECE) **(A)**, chicken embryo fibroblasts (CEF) **(B)**, baby hamster kidney cells (BHK-21) **(C)**, or bovine kidney cells (MDBK) **(D)** were infected with rNDV or rNDV_G_RABV_ (moi 0.01). Allantoic fluids and cell culture supernatants were harvested at indicated time points after infection (p. i.). Viral titers (TCID_50_/mL) were determined after titration on quail muscle (QM9) cells and subsequent immunostaining. Bar charts depict mean viral titers standard deviation ((A) n = 4, four eggs from one experiment; (B, C, D) n = 4, two samples each from two independent experiments). Asterisks indicate significant differences of mean viral titer (α = 0.05).

### Recombinant NDV_G_RABV_ expresses RABV G *in vitro* which incorporates to a limited amount into recombinant viral particles

Viral protein expression was verified by western blot analyses of DF-1 cells infected with parental rNDV and rNDV_G_RABV_. The α RABV-G serum detected RABV G with a molecular mass of ∼65 kDa, but did not react specifically with the rNDV lysates. A signal at 75 kDa was detected in all samples and was classified as an unspecific binding. NDV F_0_ (∼ 70 kDa), F_1_ (∼ 55 kDa), NP (∼ 55 kDa) and HN (∼ 70 kDa) were detected with specific primary antibodies or antisera for both recombinant viruses at their expected molecular weights (Fig. 3A). RABV G expression was additionally verified in BHK-21 and MDBK infected cells which have already proven to allow lentogenic NDV replication (Fig. 3B). Western blots of rNDV_G_RABV_ purified virions only displayed a faint band after RABV G specific antiserum incubation, indicating that a limited amount of the foreign protein is incorporated into the recombinant viral particles, whereas the NDV surface proteins F and HN were well detectable (Fig. 3C).

**Figure 3.**
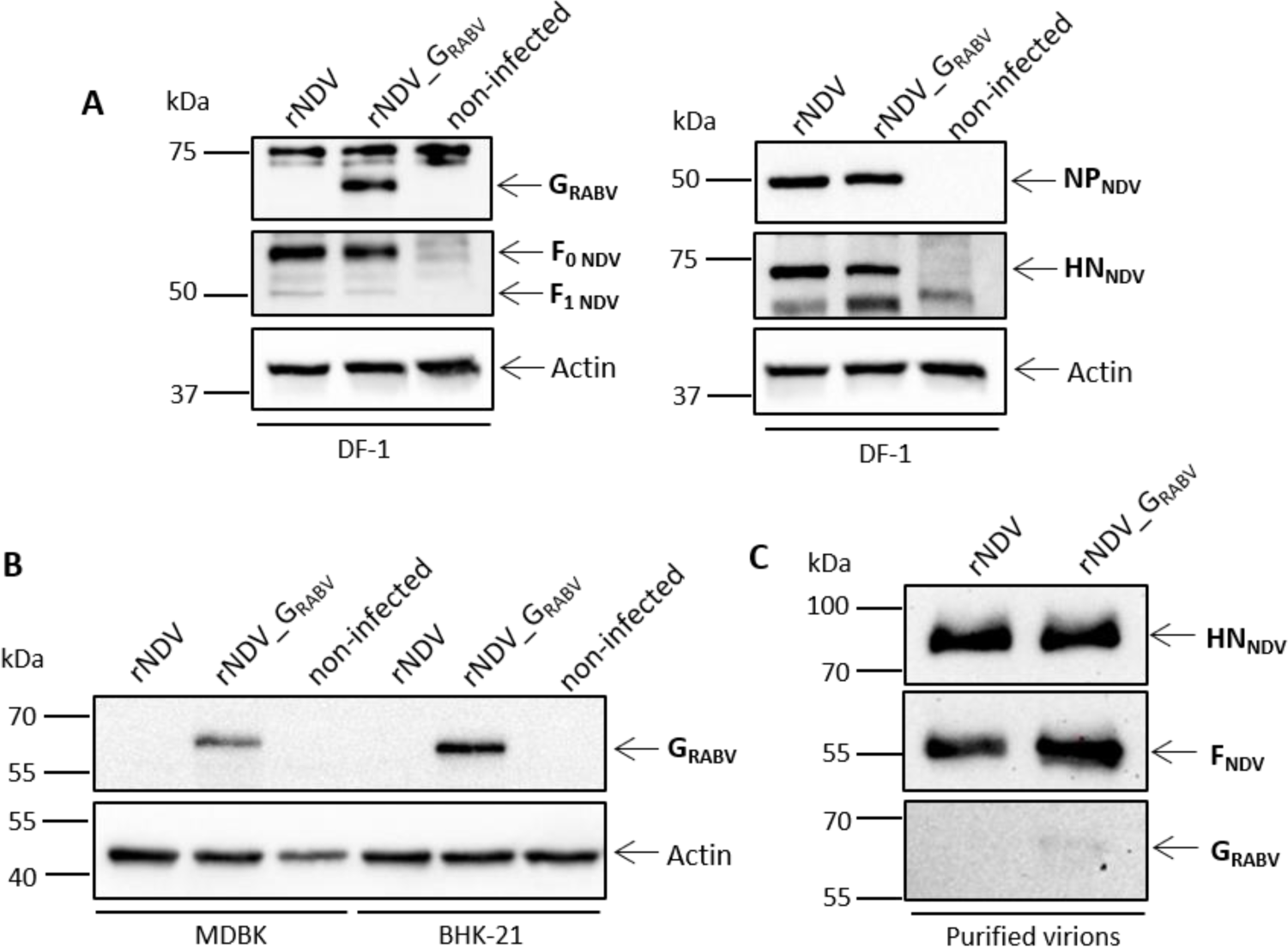
*In vitro*-analysis of viral protein expression and virion composition. Chicken embryo fibroblasts (DF-1) **(A)**, bovine kidney cells (MDBK) and baby hamster kidney cells (BHK-21) **(B)** were infected with rNDV or rNDV_G_RABV_ (multiplicity of infection (moi) 5) and harvested 24 h post infection (p. i.). Cell lysates and lysates of purified virions (10 µg per lane) **(C)** were subjected to SDS-PAGE and subsequently to Western blot analysis. Viral proteins were visualized by immunostaining with respective antibodies. Β-Actin was detected on every cell lysate blot as loading control.

Furthermore, immunofluorescence staining of rNDV_G_RABV_ infected permeabilized and non-permeabilized QM9 cells using a RABV G specific mab displayed a specific staining that is lacking in rNDV infected QM9 cells whereas NDV HN is specifically detectable in both infections (Fig. 4).

**Figure 4.**
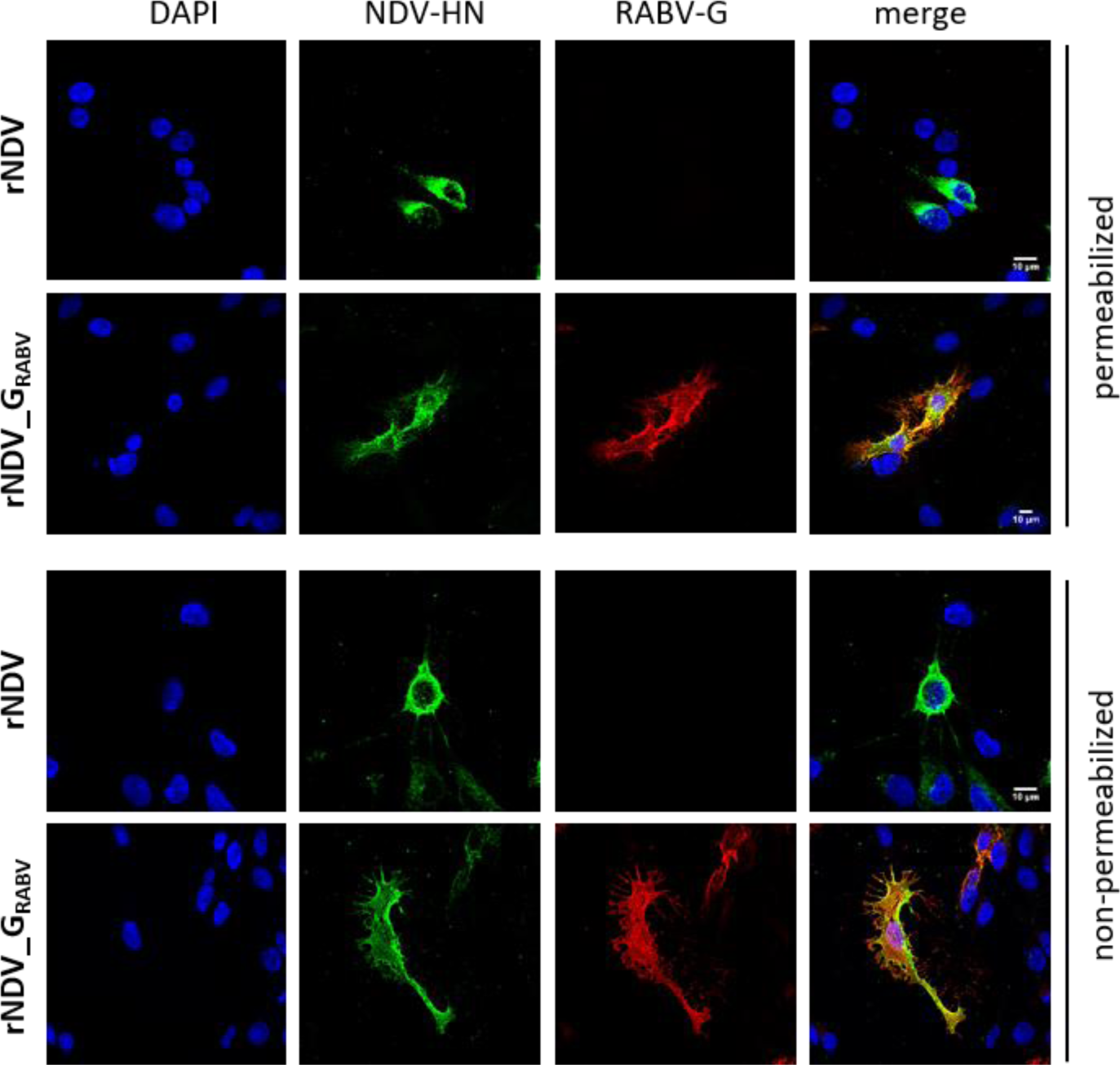
*In vitro-*analysis of viral protein expression. Quail muscle cells (QM9) were infected with rNDV or rNDV_G_RABV_ (moi 0.1), fixed 24 h p. i., optionally permeabilized with 0.1 % Triton X-100, and immunostained with respective antibodies. Cellular distribution was analyzed by confocal microscopy.

### Insertion of RABV G does not increase NDV pathogenicity in vivo

The MDT in embryonated SPF chicken eggs and the ICPI in one-day-old SPF chickens was determined to assess whether insertion of RABV G into the NDV backbone alters the *in vivo*-pathogenicity of NDV in its host species. Both ICPI and MDT suggest that the transgene insertion leads to an even higher attenuated phenotype and classifies the experimental vaccine virus as lentogenic, which is an important prerequisite of NDV based live vaccine viruses (Table 2).

**Table 2:**
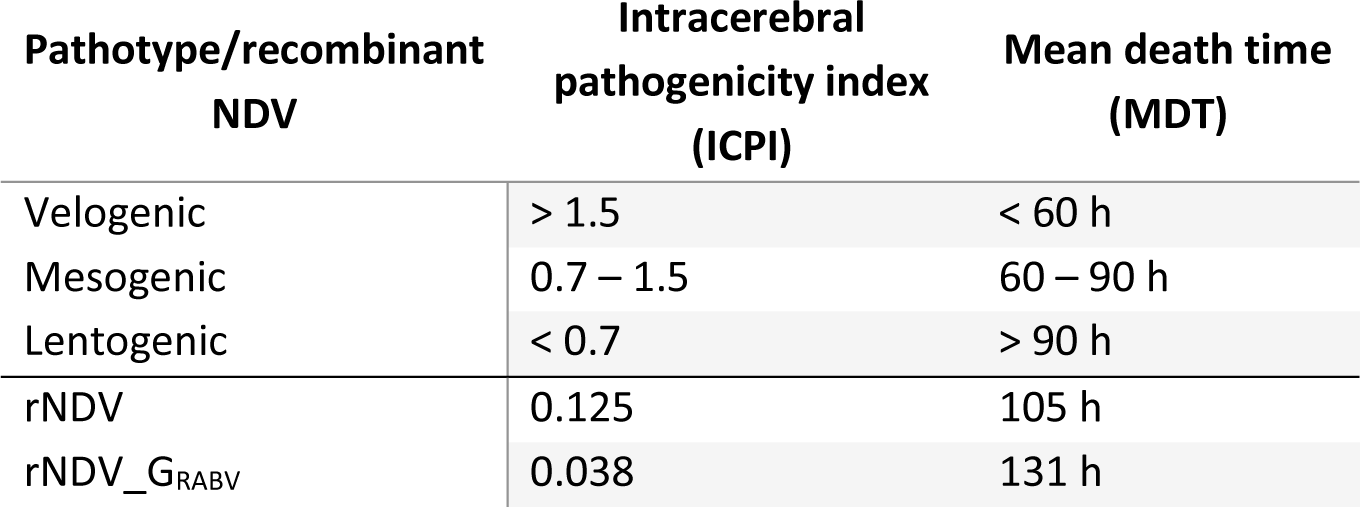
Pathotyping of rNDV and rNDV_G_RABV_ according to pathogenicity indices *in vivo*.

### Orally applicated live rNDV_G_RABV_ is safe in foxes and goats

Neither of the six goats and six foxes administered a high dose (3x10^8.75^ TCID_50_/animal) of live recombinant rNDV_G_RABV_ via the oral route, nor the control animals, which had received a similar dose of live recombinant parental rNDV showed clinical signs or altered behavior nor developed persistent fever over the period of 28 days (Fig. S1A, B). The body weight of vaccinated foxes was monitored which stayed stable during the trial (Fig. S1C).

NDV specific viral RNA was detected in swabs from both species beginning 2 dpv. Notably, the number of positive swabs and amount of detected RNA was higher in swabs taken from foxes than from goats (Fig. 5). Foxes displayed positive oral swabs until 7 dpv and goats until 4 dpv, indicating a certain degree of replication of parental rNDV and the recombinant vaccine virus in the oral cavity. One goat and four foxes shed nasally and one goat displayed a positive rectal swab. Animals that shed nasally did also shed orally with one exception (Table S1). Infectious virus could not be re-isolated in ECE from any of the positive swabs.

**Figure 5.**
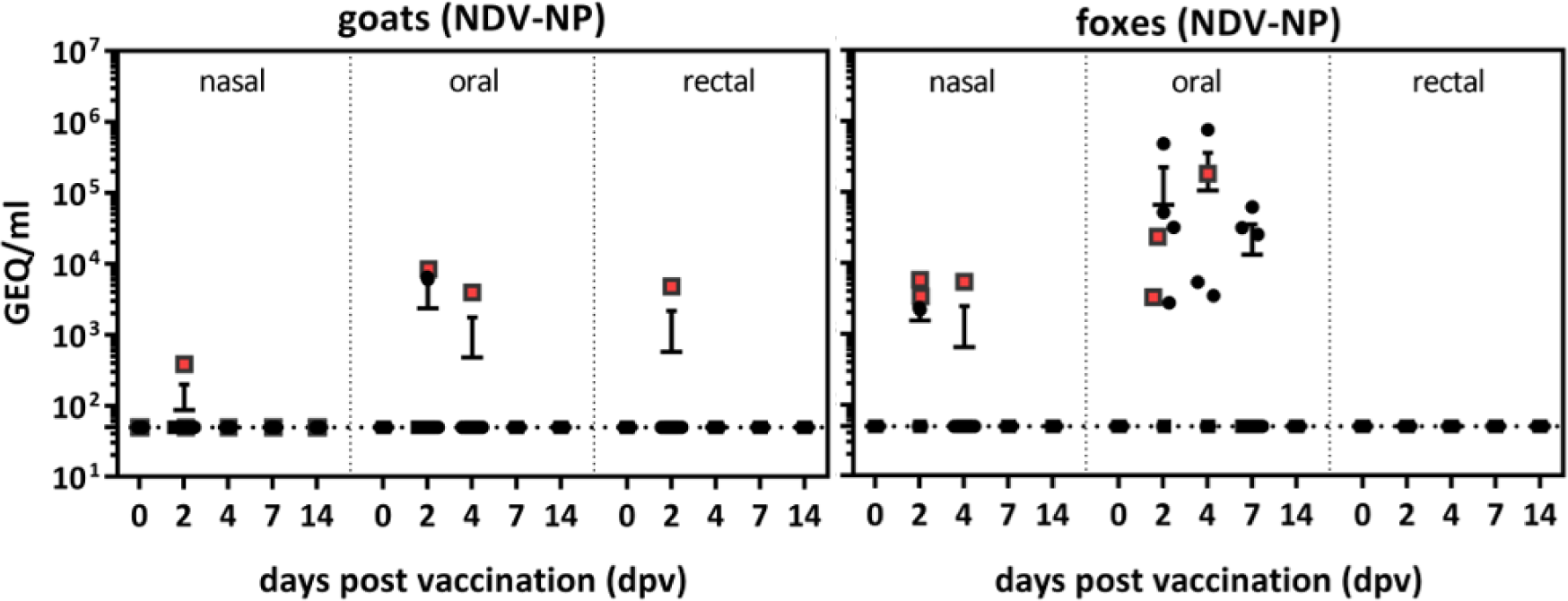
Analysis of virus replication and shedding after oral vaccination. Goats and foxes were directly orally vaccinated with either parental rNDV (n=3) or RABV G expressing rNDV_G_RABV_ (n=6). Nasal, oral and rectal swabs were taken from all animals at indicated days after vaccination (dpv) and analyzed by quantitative real-time RT-PCR (RT-qPCR) for the presence of NDV NP specific RNA. Red dots represent rNDV inoculated animals and black dots represent rNDV_G_RABV_ inoculated animals respectively.

### Orally applicated live rNDV_G_RABV_ induces NDV and RABV specific neutralizing antibodies in foxes and goats

All animals were tested NDV and RABV seronegative prior to oral vaccination (0 dpv). Vector and insert specific seroconversion was observed in both species beginning from 14 dpv onwards (Fig. 6 and 7, Table S2).

**Figure 6.**
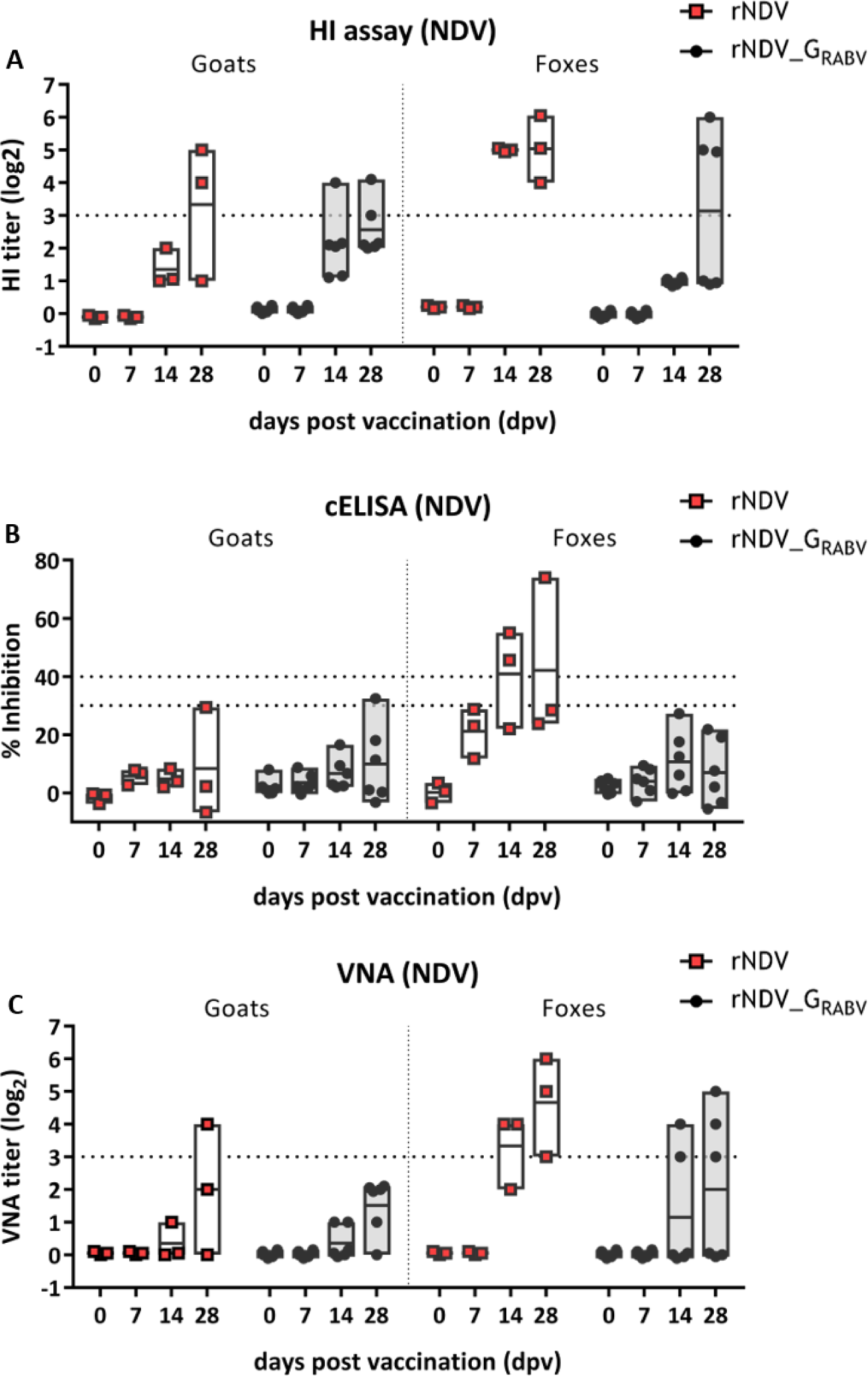
Analysis of serum antibodies against NDV after oral vaccination. Serum was taken from all animals at indicated days after vaccination (dpv) and analyzed for antibodies specific to NDV by a competitive ELISA (cELISA; seropositivity: inhibition ≥ 40 %) **(A)**, the hemagglutination inhibition (HI) assay (seropositivity: log2 ≥ 3) **(B)** and the virus neutralization assay (VNA; seropositivity: log2 ≥ 3) **(C)**. Dotted lines indicate the respective thresholds. Floating bars depict the mean titers and individual values.

**Figure 7.**
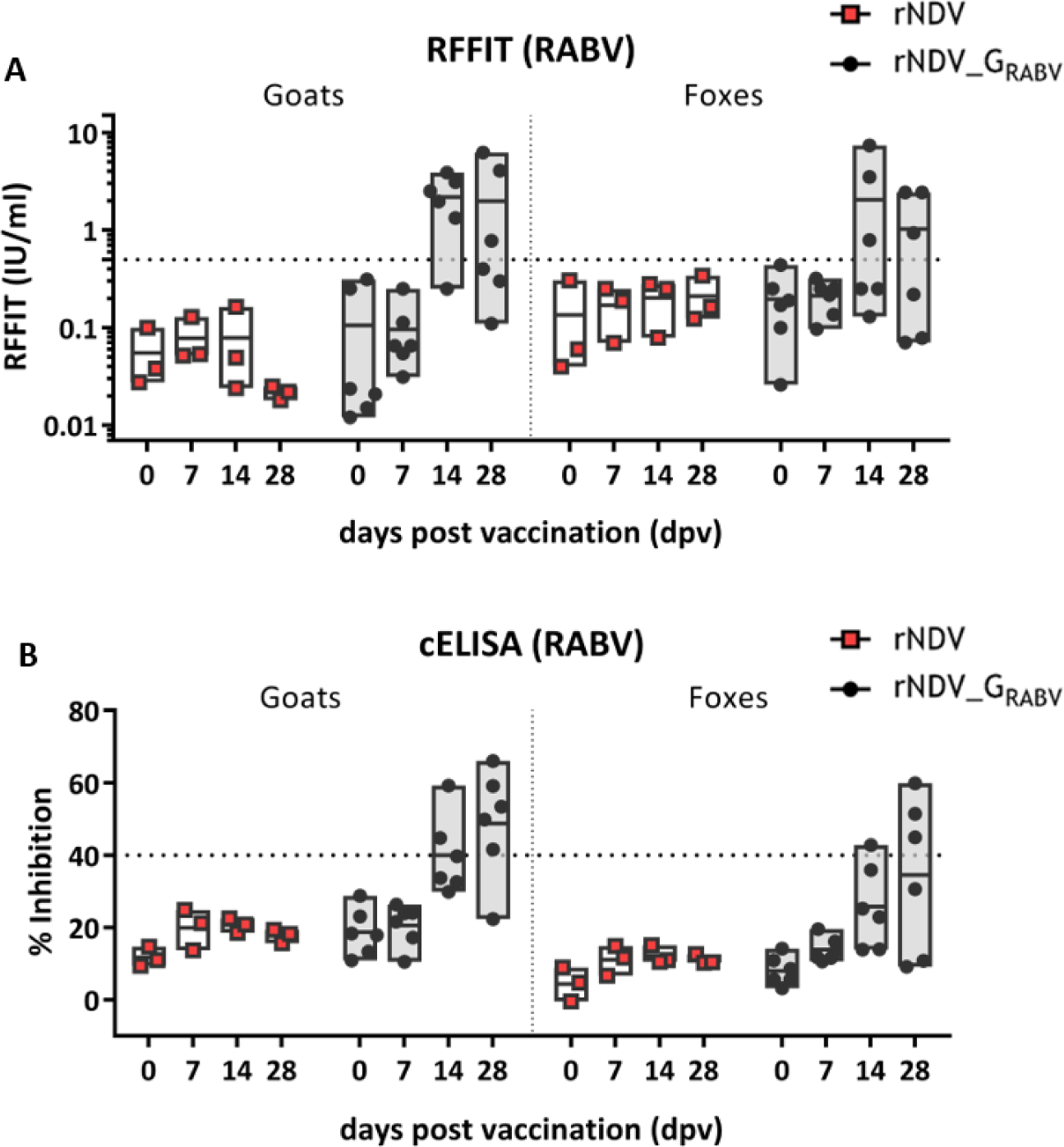
Analysis of serum antibodies against RABV G after oral vaccination. Serum was taken from all animals at indicated days after vaccination (dpv) and analyzed for RABV G specific antibodies by a competitive ELISA (cELISA; seropositivity: inhibition ≥ 40 %) **(A)** and the rabies virus fluorescent focus inhibition test (RFFIT; seropositivity: IU/mL ≥ 0.5 %) **(B)**. Dotted lines indicate the respective thresholds. Floating bars depict the mean titers and individual samples.

Two rNDV and two rNDV_G_RABV_ administered goats were tested NDV antibody positive 28 dpv by HI assay, but not by NDV ELISA. Only two of them showed values of inhibition between 30 and 40 % and were considered indeterminate (Fig. 6A, B). While all of the rNDV inoculated foxes and three out of six rNDV_G_RABV_ inoculated foxes developed NDV specific antibodies as detected by HI assay 28 dpv, only two control foxes were detected NDV seropositive by ELISA (Fig. 6A, B).

All but one rNDV_G_RABV_ vaccinated goats were tested RABV antibody positive either in ELISA or RFFIT 14 or 28 dpv and three out of six rNDV_G_RABV_ vaccinated foxes had RABV binding and neutralizing antibodies 28 dpv, while all rNDV inoculated goats and foxes remained seronegative for RABV (Fig. 7A, B).

A significant positive correlation (p < 0.0001) was observed for the NDV-ELISA and HI assay in determining NDV specific antibodies as well as for RABV-ELISA and RFFIT in detecting RABV specific antibodies in both animal species (Fig. S2).

Whereas all rNDV_G_RABV_ vaccinated foxes that seroconverted developed NDV as well as RABV specific antibodies, more rNDV_G_RABV_ vaccinated goats were tested RABV-G seropositive than NDV seropositive 28 dpv (Table 3). To verify this lack of vector immunity in those animals, a third serological assay was performed to detect NDV neutralizing antibodies by a classical VNA. Only one of the control goats displayed a distinct positive neutralizing titer whereas the other goat sera only displayed low or no neutralizing activity. NDV seropositive fox sera also exposed NDV neutralizing antibodies (Fig. 6C, Table 3).

**Table 3:**
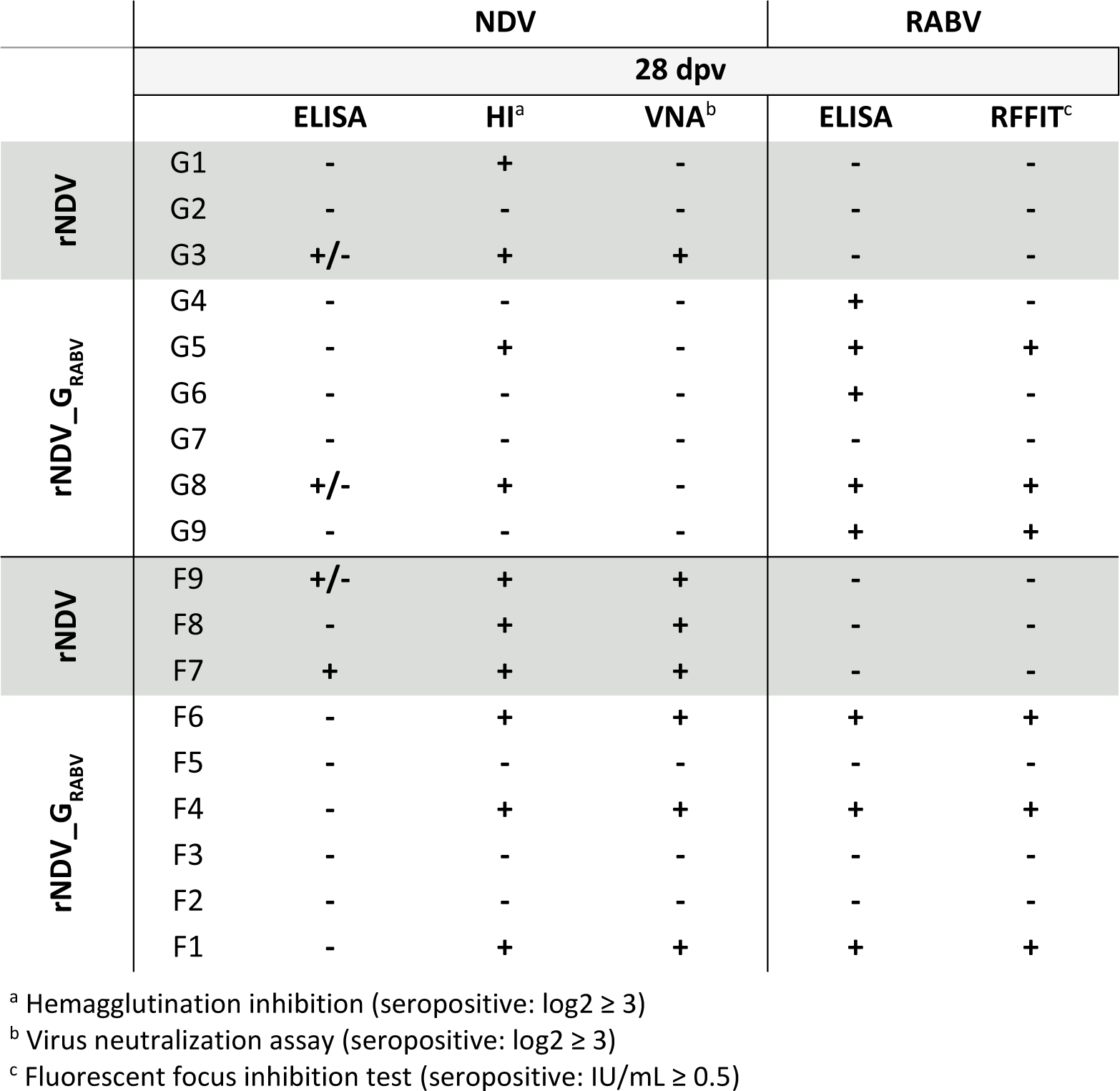
Individual outcome of NDV and RABV serological assays of goat (G) and fox (F) sera 28 days after oral vaccination (dpv) with either rNDV or rNDV_G_RABV_.

It was noticed, that primarily those animals which developed a vector or insert specific immunity also displayed NDV RNA positive swab samples after vaccination (Table S1, S2).

### Orally applicated live rNDV_G_RABV_ induced an NDV and a limited RABV specific local T cell response

The retropharyngeal lymph nodes of all animals were removed post mortem and lymphocytes isolated to investigate the local T cell mediated immune response after oral vaccination of goats and foxes. Isolated lymphocytes exhibited responsiveness after NDV and, to a lower extent, after RAVB antigen stimulation in goats and foxes. However, the number of antigen specific IFN-γ secreting lymphocytes differed greatly between individual animals. In general, foxes displayed higher numbers of antigen specific T cells than goats and more IFN-γ positive spots could be counted after stimulation with NDV than RABV (Fig. 8A). As a result, two rNDV_G_RABV_ vaccinated goats and five rNDV_G_RABV_ vaccinated foxes as well as one fox, that received rNDV, showed high numbers of IFN-γ secreting lymphocytes after NDV stimulation, whereas only one rNDV_G_RABV_ vaccinated goat and one rNDV_G_RABV_ vaccinated fox developed RABV specific T cells. Two other rNDV_G_RABV_ vaccinated foxes developed slightly elevated numbers of RABV specific IFN-y producing T cells (Fig. 8B). Unspecific lymphocyte activation with concanavalin A resulted in robust IFN-γ secretion (goats: 697 IFN-γ SFU/10^6^ cells; foxes: 576 IFN-γ SFU/10^6^ cells), indicating general responsiveness of the isolated lymphocytes, whereas none or only few spots could be observed after incubation with medium.

**Figure 8.**
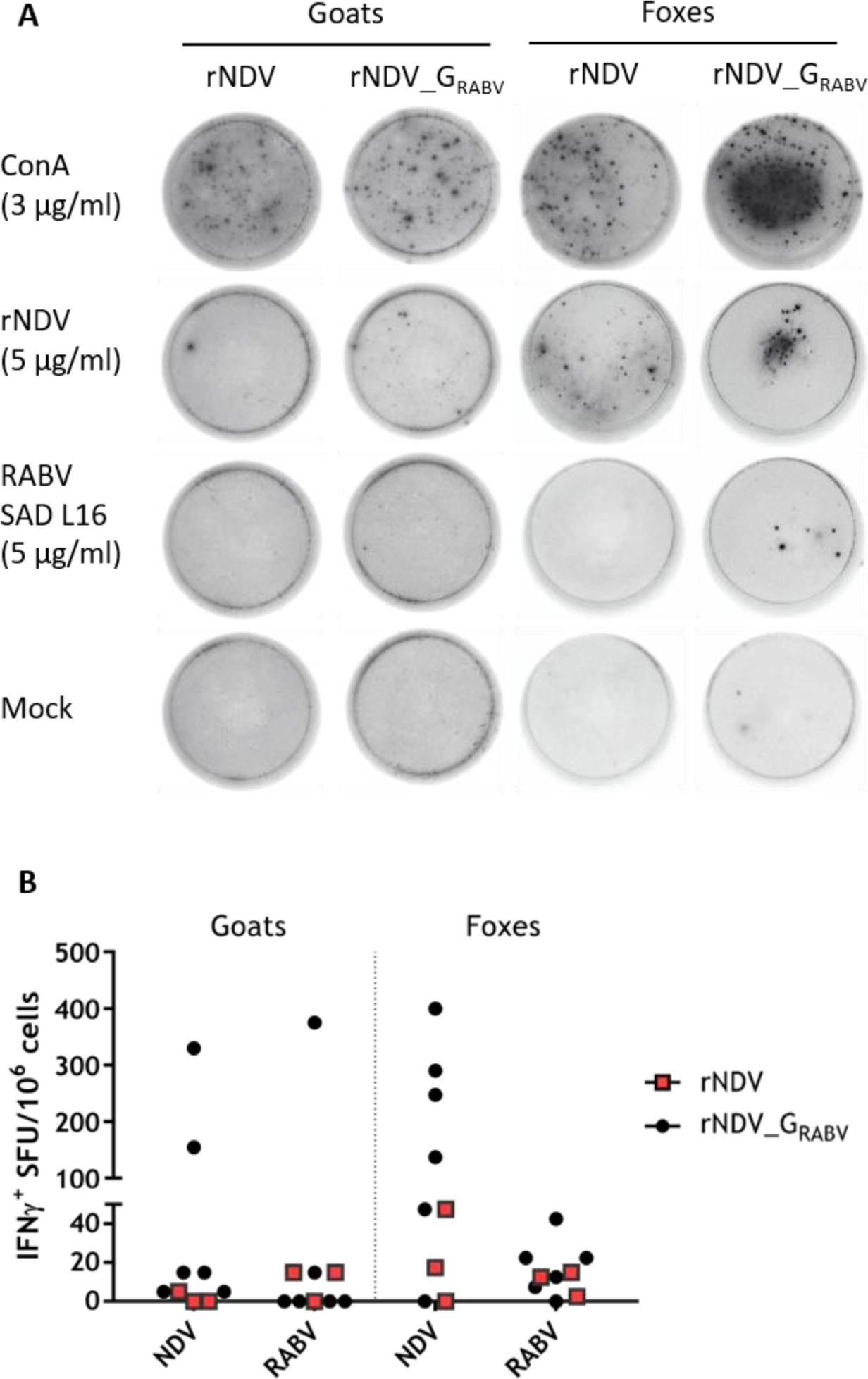
Analysis of T-cell specific IFN-γ production of pharyngeal lymphocytes from goats and foxes after oral vaccination. Pharyngeal lymph nodes from all animals were removed post mortem (28 days after vaccination). Lymphocytes were isolated and their specific interferon γ (IFN-γ) response was measured by ELISpot assays. (A) Representative images of cavities showing IFN-γ spot forming units (SFU) of Concanavalin A (ConA, positive control), medium (negative control) and NDV or RABV antigen-stimulated fox and goat lymphocytes for rNDV and rNDV_G_RABV_ vaccinated animals. (B) Graph shows antigen specific SFU per 10^6^ cells, corrected for mock control.

The IFN-γ secretion did not show any correlation with the NDV or RABV G antibody titers obtained in the sera of vaccinated goats and foxes.

## Discussion

In this immunogenicity study in goats and foxes as representatives for carnivore and herbivore species, we could demonstrate that a newly generated NDV based recombinant vector virus expressing the RABV G is safe and elicits an immune response after a single oral application (Fig. 6, 7). While a similar NDV vectored rabies G recombinant proved to be immunogenic after parenteral injection in mice, cats and dogs (8, 22), here, we demonstrated for the first time the potential of NDV as an oral vaccine vector for rabies control.

During the study, all control animals remained RABV seronegative while three foxes and five goats vaccinated with rNDV_G_RABV_ developed RABV specific antibodies as measured in ELISA (Table 3, Fig. 7). The RFFIT confirmed these results, showing a few samples that were ELISA positive but RFFIT negative. This pattern of higher positivity in ELISA as compared to RFFIT is similar to other oral vaccination studies in rabies reservoir species using modified live viruses (MLV) (37, 42, 43). Interestingly, it was shown that the ELISA is a better predictor for survival after challenge than the RFFIT (40). Against this background, it can be assumed that all ELISA positive animals would have been protected after RABV challenge infection.

In our study, we also tested for the immune response against the vector virus. In both species, VNA and HI assays were more sensitive in detecting NDV specific serum antibodies than the commercial competitive ELISA (Fig. 6), possibly because the threshold for positivity was not adjusted to mammalian sera. Furthermore, more foxes than goats developed NDV specific HI and binding antibodies, which at the same time showed higher titers, that might correlate with the prolonged shedding period of foxes compared to goats.

Interestingly, while almost all control animals which were given the parental rNDV vector developed measurable antibodies against NDV, only half of the rNDV_G_RABV_ vaccinated foxes and two out of six rNDV_G_RABV_ vaccinated goats had NDV specific antibodies. Additionally, the three rNDV_G_RABV_ vaccinated seropositive foxes displayed RABV and NDV specific antibodies, whereas some RABV antibody positive goats were tested NDV seronegative in both HI assay and ELISA (Fig. 6, Table 3). This suggests a higher immunogenicity of RABV G as compared to vector proteins. Similar observations were made in studies using rabies recombinant vaccines based on bovine herpesvirus (BHV, (44)) and canine distemper virus (CDV (45)).

In the process of virus clearance, the cell mediated immune response plays an important role in vaccine elicited protection (46). NDV is known to efficiently induce Th1 mediated cellular immunity, resulting in IFN-γ production in chickens as well as mammals (22, 47, 48). Additionally, oral vaccination of foxes using rabies vaccine strains resulted in the specific priming of PBMC, indicating a systemic T cell immune response (43, 49). Here, we could detect NDV specific IFN-γ producing regional lymphocytes from the pharyngeal lymph nodes in two goats and six foxes, indicating the activation of the cell-mediated immune system in the region of vaccine application in those animals (Fig. 8). Interestingly, although foxes showed a higher percentage of NDV serum seropositivity after vaccination, no correlation was found between individual foxes and goats that developed a humoral or cell mediated immune response, respectively. The number of RABV specific IFN-y positive lymphocytes was lower than after NDV stimulation, and fewer animals developed a T cell response. In contrast to the NDV proteins which are all expressed from the NDV vector and could contain T-cell epitopes, the spectrum of potential T cell reactions to RABV is here reduced to epitopes in the viral glycoprotein, which may explain the lower T-cell response against RABV antigens, as it has been shown that also RABV nucleoprotein displays many T cell epitopes, which are not expressed by the NDV vectored vaccine (50–52).

One of the main considerations besides immunogenicity and efficacy, are safety aspects before developing and registering an MLV vaccine construct. No goat or fox displayed any clinical signs after direct oral administration, confirming the high safety profile of NDV vectored recombinants in mammals (53–55). As shown before for other viral transgenes (8, 54, 56, 57), the insertion of the foreign gene did not alter the pathogenicity of the recombinant virus in the NDV host species (Table 2), even though few amounts of RABV G was detected in purified rNDV_G_RABV_ (Fig. 3C). Genetic stability is another criterion for safety considerations (58). For our NDV construct, this was confirmed after passaging in ECE without any mutations in the inserted RABV G or the F proteolytic cleavage site, the main NDV virulence determinant.

After a single oral application of rNDV and RABV G expressing rNDV_G_RABV_, NDV specific RNA was detectable over a period of several days in oral and nasal swabs of the vaccinated goats and foxes, indicating limited vaccine virus replication and dissemination in the oral cavity and the upper respiratory tract, but no systemic spread (Fig. 5). However, infectious virus was not re-isolated, thus an entry of infectious NDV into bird populations seems extremely unlikely. Where virus replication occurs is not known yet. The palatine tonsils of different carnivore species were reported to be the primary site for virus uptake and replication of RABV and also VACV based rabies vaccines (14, 59). While palatine tonsils are also present in ruminants like goats (60), it is unclear whether this lymphatic tissue represents a specific entry and replication site for NDV. In poultry, epithelial cells of the respiratory and gastrointestinal tract represent the main target cells for NDV, and as NDV binds to sialic acid containing cellular molecules, the virus can infect a variety of different avian and mammalian derived cell types, as was shown in cell culture (Fig. 2).

Both the number of positive samples, the duration of shedding and the viral load was higher in foxes than in goats (Fig. 5), indicating a higher level of susceptibility to NDV. However, it might also be related to the slight differences in the virus application process. Foxes had to be sedated for safety reasons. Therefore, the virus containing fluid may have been longer in contact with the tongue mucosa while the virus was directly applied to the oral cavity of goats and immediately swallowed. Also, goats were allowed to directly eat after virus administration.

## Conclusion

The fact that a single oral administration of a live NDV vectored rabies vaccine is able to induce a systemic humoral and a local cell mediated immune response in foxes and goats opens new avenues for vaccine development. In this proof-of-concept study with a limited number of animals, an immune response was detected in some but not all animals, clearly indicating the need for improvements if the requirements for registrations by EMA (58) and WOAH (61) are to be met. Potential parameters for an improved immune response are genetic modifications to the gene insert to increase its stability (62). Also, in this study, a dose of 10^8.5^ TCID_50_/animal was used. However, NDV can replicate to even higher titers in embryonated chicken eggs, and data from RABV MLV suggest a dose-response correlation. Generally, rabies in dogs and e.g. kudus is a public health burden particularly in low- and middle-income countries (LMICs) where investments into disease control are largely hampered by insufficient funds. With NDV as a vector, vaccines can be manufactured similarly to influenza virus vaccines at low cost in embryonated chicken eggs in facilities located globally, including in LMICs. Beyond rabies, it is likely that NDV based vaccines generally elicit an immune response after oral delivery, thus giving hope to vaccines for animal diseases where parenteral vaccinations face limitations, particularly in the outreach to remote communities in LMICs.

## Acknowledgments

The authors thank Martina Lange and Jeannette Kliemt for excellent technical assistance, as well as Christine Luttermann and Kristin Trippler for sequencing services during the generation and *in vitro* characterization of the recombinant virus. The authors thank all animal care takers of the Friedrich-Loeffler-Institute for their fostering of animals and their support during sampling and Alexander Schäfer for helpful discussion and support regarding the lymphocyte preparation and IFN-γ ELISpot assay.

## Author Contributions

Conceptualization: SF, TCM, ARO, CF, TM, MM; Data curation: MM, CF, TM, DPB; Formal analysis: MM, CF; Investigation: MM, DPB, CF, TM, CG; Supervision: TCM, ARO, SF; Visualization: MM; Writing – Original Draft Preparation: MM, CF, TM; Writing – Review & Editing: all authors

## Supporting information

**Supplemental figure 1. Development of temperature and weight after direct oral immunization. (A)** Goats and **(B)** foxes were directly orally vaccinated with either parental rNDV (n=3) or RABV G expressing rNDV_G_RABV_ (n=6). Rectal temperature of goats and foxes as well as weight of foxes was monitored at indicated timepoints after oral vaccination.

**Supplemental Figure 2. Correlation between NDV binding antibodies (ELISA) and hemagglutination inhibition antibodies (HI assay) or between RABV binding antibodies (ELISA) and RABV neutralizing antibodies (RFFIT) in oral vaccinated goats and foxes. (A)** Goats and **(B)** foxes were directly orally vaccinated with either parental rNDV (n=3) or RABV G expressing rNDV_G_RABV_ (n=6). Sera of all animals was tested for NDV and RABV G specific antibodies at different timepoints after vaccination as described in material and methods section. Pearson correlation coefficient [-1; +1] was calculated to determine extent of correlation between serological assays in determining NDV or RABV specific antibodies.

**Supplemental table 2. Individual serological data after oral vaccination of goats and foxes.** Goats and foxes were directly orally vaccinated with either parental rNDV (n=3) or RABV G expressing rNDV_G_RABV_ (n=6). Serum was taken from all animals at indicated days after vaccination (dpv) and analyzed for antibodies specific to RABV by a competitive ELISA (cELISA; seropositivity: inhibition ≥ 40 %) and the fluorescent focus inhibition test (RFFIT; seropositivity: IU/mL ≥ 0.5). Serum was analyzed for antibodies specific to NDV by a competitive ELISA (cELISA; seropositivity: inhibition ≥ 40 %) and the hemagglutination inhibition (HI) assay (seropositivity: log2 ≥ 3). Seropositive samples are highlighted in green, indeterminate samples in orange.

